# The plant specific cohesin subunit SYN4 contributes to 3D genome organization

**DOI:** 10.1101/2024.03.27.586521

**Authors:** Pirita Paajanen, Carsten Proksch, Sylvia Krüger, Karin Gorzolka, Jörg Ziegler, Tina Romeis, Antony N. Dodd, Kirsten Bomblies, Vinzenz Handrick

## Abstract

Chromatin architecture in the cells of animals and fungi influences gene expression. The molecular factors that influence higher genome architecture in plants and their effects on gene expression remain unknown. Cohesin complexes, conserved in eukaryotes, are essential factors in genome structuring. Here, we investigated the relevance of the plant-specific somatic cohesin subunit SYN4 for chromatin organisation in *Arabidopsis thaliana*.
Plants mutated in *SYN4* were studied using HRM, Hi-C, RNA sequencing, untargeted and targeted metabolomics and physiological assays to understand the role of this plant-specific cohesin.
We show that *syn4* mutants exhibit altered intra- and interchromosomal interactions, expressed as sharply reduced contacts between telomeres and chromosome arms but not between centromeres, and differences in the placement and number of topologically associated domains (TADs)-like structures. By transcriptome sequencing, we also show that *syn4* mutants have altered gene expression, including numerous genes that control abiotic stress responses.
The response to drought stress in *Arabidopsis* is strongly influenced by the genome structure in *syn4* mutants, potentially due to altered expression of *CYP707A3*, an ABA 8’-hydroxylase.

**In brief:** The 3D architecture of the genome extensively influences gene expression in animals and fungi. We show that the plant-specific cohesin subunit SYN4 affects intra- and interchromosomal interactions including telomere clustering, with consequences for the expression of genes of transient and induced biological pathways and the biosynthesis of bioactive compounds. Highly condensed genome structures at the *CYP707A3* locus positively affects the stress response to water deprivation regulated by abscisic acid.

## Introduction

The geometric arrangement and packaging of chromatin plays an important role in controlling gene expression (Merkenschlager and Nora, 2016). A major player in chromatin packaging in animal, fungi and bacteria is the cohesin complex, which is essential for sister chromatid cohesion and compacting and organizing chromatin (Nasmyth and Haering, 2009). During interphase of somatic cells, cohesins shape the higher genome architecture through a so-called “loop extrusion”, in which DNA is actively pulled in an ATP-dependent manner through a ring formed by the cohesin complex (Kim et al., 2019; Mizuguchi et al., 2014). The resulting chromatin loops form the building blocks for higher-order topologically associated domains (TADs), which play key roles in gene regulation (Greenwald et al., 2019; Xiao et al., 2021).

While many cohesin functions are known to be conserved between plants and animals (Peters et al., 2008), whether plant cohesins also play a role in managing 3D genome architecture is unknown. All cohesin complexes include STRUCTURAL MAINTENANCE OF CHROMOSOMES proteins, SMC1 and SMC3, and several other proteins that form a loop surrounding the DNA (Fig. 1a; Kim et al., 2019; Davidson et al., 2019). These proteins are then linked at their bases by an additional component, an α-kleisin, that varies among functionally distinct cohesins. The main functional difference between meiotic and the mitotic cohesin complexes (which differ in their chromosome-association dynamics) arises from the α-kleisin component: Meiotic cohesin contains the α-kleisin REC8, while the mitotic complex contains the α-kleisin RAD21/SCC1 (Fig. 1a; Nasmyth and Haering, 2009). In *A. thaliana*, there are four different α-kleisins (SYN1/REC8; SYN2/RAD21.1; SYN3/RAD21.2; SYN4/RAD21.3; Fig. 1a). SYN1 is purely meiotic, and *syn1* mutants are sterile and show a range of meiotic defects in recombination, axis formation, synapsis and chromosome segregation (Bai et al., 1999; Bhatt et al., 1999). SYN3 is similar in sequence to SYN1, but unlike SYN1, is essential for viability, and is present in both somatic and meiotic cells (Jiang et al., 2007). SYN2 and SYN4 seem to be homologs of RAD21, the mitotic α-kleisin of yeast (da Costa-Nunes et al., 2006 and 2014). Like RAD21, both proteins appear to have mitotic functions, with SYN2 playing a role in DNA repair (da Costa-Nunes et al., 2006; Dong et al., 2001) while SYN4 plays a role in genome stability and is associated with replication factors (da Costa-Nunes et al., 2014). Whether or how these different mitotic cohesins might affect chromatin architecture is as yet unknown.

**Fig. 1.**
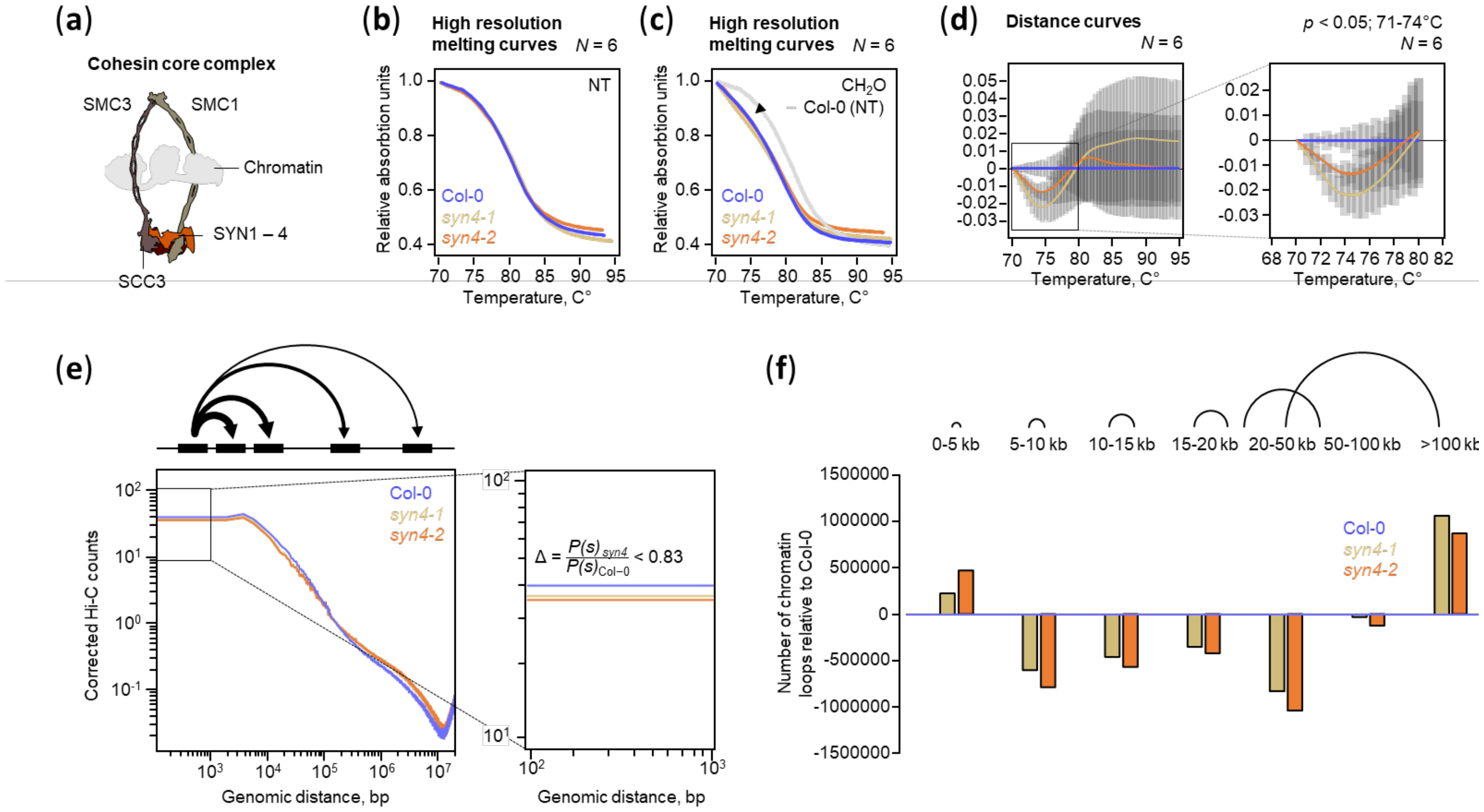
Analy ses of the DNA interactom. **(a)** Cartoon of the cohesin core complex that has bound clironiatin. The cohesin core complex contains two SMC (STRUCTURAL MAINTANANCE OF CHROMOSOME) arms. SMC1 and SMC3. which are linked by α-kleisins. Four α-kleisins exist in Aiabidopsis (SYN1-4). SCC3 mediates the topology-specific binding of the cohesin complex. **(b-c)** Insertions in *SYN4* effect DNA thermostability. Normalized melting profiles from six independent HRM (high resolution melt) experiments without (no treatment = NT. 2A) and with formaldehyde-fixed genomic DNA (CH2O. 2B) from manire leaves. **(d)** Distance curves, with Col-0 as reference, show dial DNA of the *syn4* mutants starts to dissociate at lower temperatures, suggesting lower interactions in die fixed DNA (paire-wise Student’s T-test between Col-0 and *syn4*-l respectively Col-0 and syn4-2 measurements). Grey bars represent standard errors, s.e. **(e)** Genome-wide Hi-C counts versus genomic distance (P(s)) in Col-0 wild type (blue). syn4-1 (yellow) and syn4-2 (orange). **(f)** A high number of Col-0 clironiatin loops larger than 5 kb and smaller than 50 kb are no longer formed in the *syn4* mutants.

Here, we sought to test whether cohesin might also play a role in genome architecture in plants. Because SYN3 is essential (Jiang et al., 2007), and thus no knock-out mutant is available, SYN1 is meiosis-specific (Bai et al., 1999, Bhatt et al., 1999), and SYN2 has been reported to function specifically in DNA repair (da Costa-Nunes et al., 2006; Dong et al., 2001) we focused our efforts on the remaining mitotic α-kleisin, SYN4/RAD21.3. SYN4 physically interacts with chromatin remodelers that were not found for the other α-kleisins (McWhite et al., 2020; BioGrid: https://thebiogrid.org/16763/summary/arabidopsis-thaliana/syn4.html), so represents a good candidate for investigation. We identify that SYN4 contributes to 3D genome architecture, and this alters gene expression and plant physiology. Therefore, our results suggest that plant cohesins have a role that is conserved with cohesins from other eukaryotes, in regulating 3D chromatin architecture via both inter- and intra-chromosomal interactions. Furthermore, our findings indicate that defined genome structures are needed to respond to changing environmental influences.

## Materials and Methods

### Plant material and cultivation

The α-kleisin T-DNA insertion mutants (*syn4-*1; SALK_020171, *syn4-*2; Schubert et al., 2009) were obtained from the Nottingham Arabidopsis Stock Centre (http://arabidopsis.info/). Insertion lines were genotyped with primers described in Schubert et al. (Schubert et al., 2009) and homozygous lines were generated. Seedlings were grown under short-day conditions (8h light, 16h dark) with 120 μmol·m-2s-1 light intensity, with temperatures of 22L□°C (light period) and 20L□°C (dark period).

### Fixation of leaf material using formaldehyde and extraction of genomic DNA

The plant material of all genotypes was cut with a scalpel into pieces of about 1x1 cm and put into a beaker with a stirring fish and poured on with fixing buffer (20 mM Hepes pH 8.0, 250 mM sucrose, 1 mM MgCl_2_, 5 mM KCl; 40% (v/v) glycerol, 0.25% (v/v) Triton X-100; 2% (v/v) formaldehyde, 0.1 mM β-mercaptoethanol and 0.1 mM PMSF). The plant material was stirred at high rpm until it sank (approx. 30 min). The fixation was stopped by adding glycine (1 M, final concentration) while stirring for 5 min. The plant material was then washed 6 times with ice-cold sterile double-distilled water. Finally, the plant material was dried on paper towels and shock frozen. For high resolution DNA melt analysis genomic DNA was extracted from frozen fixed and non-fixed material according to a published method (Edwards et al., 1991).

### High resolution DNA melt analysis

2000ng DNA (fixed or non-fixed) was mixed with 10μL Brilliant III SYBR Green qPCR Master Mix (Promega) and the DNA melting behaviour was analysed in the CFX96 Touch Real-Time PCR Detection System (Bio-Rad). Samples were conditioned at 25°C for 5 minutes and heated from 70°C, at 0.5°C intervals every 30 seconds, to 95°C. The resulting melting curves were exported from the CFX96 system.

### Sample preparation and sequencing for chromatin capture analyses

Sample preparation and sequencing for chromatin capture analyses were contracted to Phase Genomics (Seattle, Washington, USA). Fixed plant material was shipped on dry ice. We received the FASTQ raw data for the bioinformatic analyses.

### Hi-C heat map

The raw fastq sequences were downloaded from Phase Genomics (Seattle, Washington, USA) and checked. The Hi-C reads were processed by juicer v1.6 (Durand et al., 2016), and HiC contact matrices were produced. The restriction enzyme was DpnII, and TAIR10 version of the Arabidopsis thaliana was used.

### Analysis and visualisation of interchromosomal interactions

Hi-C heat maps were displayed with Juicebox 1.11.08 and the maps were normalized based on the coverage (Sqrt). The resolution was 250kbp and the respective interactions were displayed in grey values. Surface plots from selected interchromosome interactions were generated from the respective heat maps using imageJ. The relative interaction frequency always refers to the strongest blackening in the respective map.

### TAD calling

The TADs were called from the HiC contact matrix using juicer_tools v1.22 arrowhead command.

### Matching the TAD-like structures

To determine the correspondence of the TAD-like structures, the graphs from the wild type and the *syn4* insertion mutants were superimposed. The percentage of TAD areas that were not present in the respective graphs could thus be determined. The size of the areas was measured using ImageJ.

### Comparing TAD-like structures

For the differential TAD analyses, we used TADCompare as described in Cresswell and Dozmorov, 2020 (Cresswell et al., 2020).

### Extraction of RNA

Frozen plant material, was ground to a fine powder in a liquid nitrogen pre-cooled mortar and RNA was extracted using the RNeasy Plant Mini Kit (Qiagen) according to the manufacturer’s instructions. Nucleic acid concentration, purity, and quality were assessed using a spectrophotometer (NanoDrop 2000c; Thermo Scientific) and Agilent 2100 Bioanalyzer (Agilent Technologies).

### RNA-seq analyses

The analyses of the transcriptome were conducted by Novogene Co., Ltd (Cambridge, United Kingdom). Extracted total RNA was shipped on dry ice. For transcription data analysis, adapters, poly-N sequences and low-quality reads were removed from the raw data. Paired-end reads were mapped to the Arabidopsis thaliana reference genome (https://plants.ensembl.org/Arabidopsis_thaliana/Info/Index) using HISAT2. For quantification of gene expression, featureCounts was used to obtain RPKM values (Reads Per Kilobase of exon model per million mapped reads). DESeq2 yielded the differential gene expression between genotypes (three biological replicates per condition). The resulting P-values were adjusted using Benjamini and Hochberg’s approach to control for false discovery rate (FDR). Genes with an adjusted P-value < 0.05 found by DESeq2 were classified as differentially expressed.

### Metaboyping and phytohormone analysis

Metabolites of deep frozen leaf samples were extracted with 80% aqueous methanol. 1 μl of methanolic extracts (0.33 mg fresh weight / μl) were analysed with an Acquity UPLC (Waters) hyphenated to a micrOTOF-Q (Bruker). The chromatography gradient was: 0 to 1 min isocratic 95% A (water/formic acid, 99.9/0.1 [v/v]) and 5% B (acetonitrile/ formic acid, 99.9/0.1 [v/v]); 1−16 min linear from 5 to 95% B; isocratic 95% B to 18 min; 18.0−18.2 min to 5% B; 18.2−20 min isocratic 5% B. Eluting compounds were detected in positive ionization mode from m/z 50 to 1600. Data were processed with the T-Rex 3D algorithm (LC-Q-TOF) in MetaboScape 4.0 (Bruker Daltonics) and exported for further statistics in MetaboAnalyst 5.0 (https://www.metaboanalyst.ca/; Pang et al., 2021). The flavonoid content was annotated with internal libraries generated from authentic standards implemented in MetaboScape. The fumarate contents were analysed as described by Chutia et al., 2021.

The phytohormones analysis was carried out as described in Balcke et al., 2012.

### Transpiration assays and stomatal conductance

To measure the steady state water loss from untreated plants, 4 or 5 leaves per genotype were detached, and the continuous weight loss over time per gram of starting leaf material was determined. A calibrated SC-1 porometer (Meter Group, Inc. USA) was used to determine stomatal conductivity at the same time on each day of drought (Fig. S6b; ZT = 4). Measurements were taken from three leaves of six plants per genotype.

### Measurement of chromatin interaction with FRET probes

In the promoter region and gene body of *CYP707A3*, primer sequences were determined within frequently occurring DNA interaction sites using Geneious Prime. The primer sequences were designed having a melting temperature equal to or higher than 60°C, a GC content of between 35 and 55%, and a primer length in the range of 20 to 25 nucleotides. europhins Genomics (https://eurofinsgenomics.eu) manufactured the FRET probes (*CYP707A3*-probe-fwd, [ROX]GCAAAAGAACTTAGGGAGGTC; *CYP707A3*-probe-rev, GCATTCCATCAGTCATTACATTCGT[FLU]). The probes were diluted 1:100 with distilled water and 1μL mixed with 1500 ng of fixed or non-fixed DNA. FRET measurements were performed using a CFX96 Touch Real-Time PCR Detection System (Bio-Rad). Along a melting curve recorded between 55 and 95°C, the change in fluorescence (dR/dT) was measured. Biological replicates were analysed as duplicates. The background signals from measurements with non-fixed DNA were subtracted from the FRET signals with fixed DNA.

### Preparation of cDNA and RT-qPCR analysis

Prior to cDNA synthesis, 1 mg RNA was treated with 1 unit of DNase (Fermentas). Single-stranded cDNA was prepared from the DNase-treated RNA using GoScript Reverse Transcriptase System (Promega).

Relative quantification of gene expression was analysed by RT-qPCR using CFX96 Touch Real-Time PCR Detection System (Bio-Rad). For the amplification of gene fragments with a length of 150 to 250 bp, specific primer pairs were designed having a melting temperature equal to or higher than 60°C, a GC content of between 35 and 55%, and a primer length in the range of 20 to 25 nucleotides (*CYP707A3*-fwd, ATCCGGGGAAATTCGATCCG; *CYP707A3*-rev, GCTAGGCCCTACGATTGACC). Primer specificity was confirmed by agarose gel electrophoresis and melting point curve analysis. Primer pair efficiency was determined using the standard curve method with fivefold serial dilution of cDNA and was found to be 105%. As reference, the UBC gene was used (AT5G25760; Czechowski et al., 2005). 1 μL cDNA was used in 10-μL reactions containing Brilliant III SYBR Green qPCR Master Mix (Promega). Biological replicates were analyzed as duplicates. The PCR consisted of an initial incubation at 95°C for 3 min followed by 50 amplification cycles with 20 s at 95°C and 20 s at 60°C. For each cycle, reads were taken during the annealing and the extension step. At the end of cycling, the melting curves were gathered for 55 to 95°C. Data for the relative quantity of calibrator average (dRn) were exported from the CFX96 system.

### Protein structure analysis

The online tool PSIPRED was used to predict secondary protein structures (Jones et al., 1999; bioinf.cs.ucl.ac.uk/psipred/, last accessed February 3, 2021). The PSIPRED algorithm calculates the likelihood of local amino acid interactions including coil (C; disordered), helix (H), or sheet structures (E) for every amino acid position. The algorithm includes two feed-forward neural networks that perform an analysis of the output of PSI-Blast (position-specific iterated-Blast), which in turn is based on an alignment of multiple protein sequences. The three-dimensionally modelled protein structures of the investigated α-smallisins were downloaded from AlphaFold (https://alphafold.ebi.ac.uk/). The N-termini (1-115aa) were visualized and aligned using the PyMOL super-alignment function.

### Sequence analysis and tree reconstruction

Annotated and characterized α-kleisin protein sequences from eukaryotic species (*Arabidopsis thaliana, Aquilegia coerulea, Caenorhabditis elegans, Chlamydomonas reinhardtii, Coffea arabica, Drosophila melanogaster, Helianthus annuus, Homo sapiens, Marchantia polymorpha, Medicago truncatula, Mus musculus, Nymphaea colorata, Oryza sativa, Saccharomyces cerevisiae, Schizosaccharomyces pombe, Triticum aestivum and Zea mays*) were downloaded from the NCBI database (https://www.ncbi.nlm.nih.gov/gene). For the multiple sequence alignment, the Clustal Omega alignment algorithm implemented in Geneious Prime was used (default settings, Data S6). Based on the alignment, an unrooted phylogenetic tree was reconstructed using the Geneious Tree Builder. Default settings were adopted using the Neighbor-Joining algorithm with outgroup (Genetic Distance Model: Jukes-Cantor). A bootstrap resampling analysis with 1000 replicates was performed to evaluate the tree topology.

### Statistical Analysis

Statistical significance was tested using analysis of variance in GraphPad PRISM 5.03 for Windows (GraphPad Software). Whenever necessary, the data were *ln* transformed to meet statistical assumptions, such as normality and homogeneity of variances.

## Results

### *syn4* mutants show altered chromatin structure

In animals and fungi, a canonical function of cohesin is in the formation of chromatin loops, which form building blocks for higher 3D genome structures such as TADs (Merkenschlager and Nora, 2016). The extent of formation of these loops and domains can be assessed indirectly by assaying chromatin interactions within the genome. To test the hypothesis that SYN4 participates in the formation of chromatin loops, we investigated DNA topology in *syn4* mutants using two techniques: high resolution DNA melting analyses (HRM) and chromatin capture analyses (Hi-C).

HRM is based on observing differences in the melting behaviour of fixed double-stranded DNA in the presence of SYBR Green, which allows quantification of DNA interactions. We found that 10 and 14% of fixed genomic DNA from mature leaves of *syn4-1* and *syn4*-2, respectively, melts significantly earlier compared to wild-type DNA (Fig. 1b-d). This indicates that there are fewer cross-links, and thus chromatin interactions, in the mutants than in wild type. HRM, however, gives only a general overview, and does not provide a clear picture of the genomic distribution of either maintained or altered contacts.

To get a more precise picture of where contacts are maintained or altered in the mutant plants, we used Hi-C on pooled mature leaves of wild type Col-0 and *syn4* mutants (also in the Col-0 background). First, consistent with the HRM data, contact plots show that the two *syn4* mutants we analyzed have generally reduced contacts especially between chromatin regions that lie between 1-100 kb apart (Fig. 1e). 12-20% of the loops up to 50 kb are no longer present in the *syn4* mutants (Fig. 1f).

Hi-C further revealed striking differences in inter-chromosomal interactions. Hi-C heat maps of the Col-0 wild-type showed strong interactions of telomeres and centromeres of all chromosomes, consistent with previous findings (Grob et al., 2014, Moissiard et al., 2012), in addition to other intra-chromosome arm interactions (Fig. 2a and 2b). From the average interactions of centromeres and telomeres, a simplified map of high-frequency chromatid interactions in space can be derived and compared (Fig. 2a). Importantly, in the *syn4* mutants telomere (but not centromere) inter-chromosomal interactions are absent or strongly reduced (Fig. 2c-f, Fig. S2), which probably results in the order of chromosomes in the Hi-C heat maps being arbitrary (Fig. 2c and 2e), unlike in wild type (Fig. 2b). Therefore, it seems the telomeres and chromosome arms can move freely in the nuclear space in the *syn4* mutants, but not in wild type (Cartoons in Fig. 2a, c, e).

**Fig. 2.**
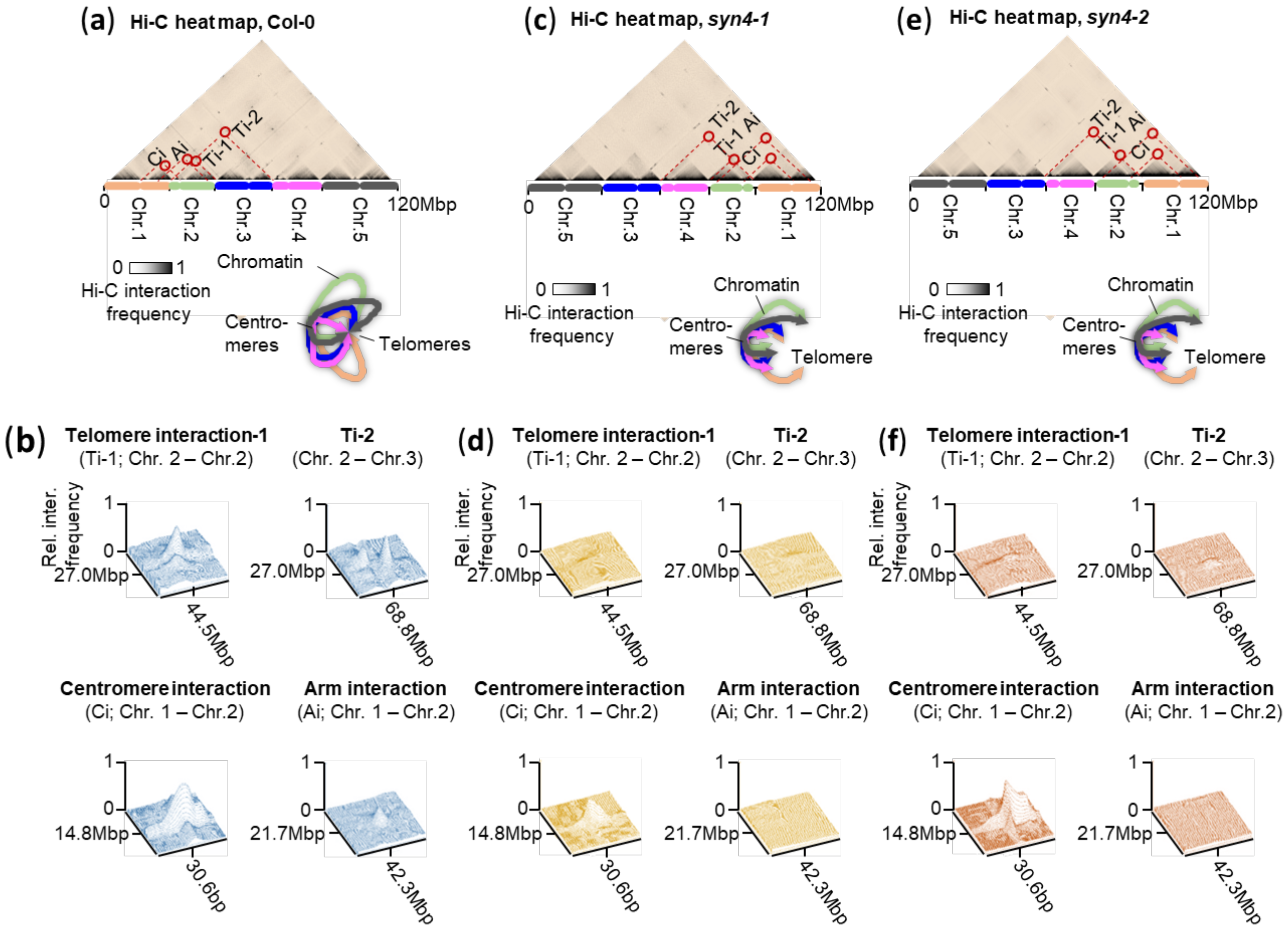
Chromatin interaction analysis. (a,c,e) Hi-C interaction heat maps generated from Col-0, *syn4*-1 and *syn4*-2 merged leaf samples. The heat maps were displayed with Juicebox 1.11.08 and the maps were nonnalized based on the coverage (Sqrt). The resolution was 250kbp and the respective interactions were displayed in grey values. Intra- and inter chromosome interaction including interaction of telomeres, centromeres and chromosome anns are indicated. Below the Hi-C heat maps, airnngement of chromatids in the nucleus based on the inter-chromosome interactions studied. (b,d,f) Smface plots show intra- and inter-chromosome interactions of chromosome arms, centromeres and telomeres generated from heat maps using imageJ. The relative interaction frequency always refers to the strongest blackening in the respective map.

Interacting domains or TAD-like regions are defined as chromosome regions with extensive interactions within but not between them. These domains tend to be less transcriptionally active, consistent with having more densely packed chromatin (Wang et al., 2015). These TAD-like regions are interspersed by transcriptionally active and open chromatin (Wang et al., 2015) and can be identified based on regions of enriched contact density in Hi-C maps. We analyzed the genome and found that in the *syn4-1* mutant, only 37% of the TAD-like structures match those of Col-0 wild type (Fig. 3a; Data S1). This was similar in *syn4*-2 and the TAD-like organization is largely consistent between the two mutant lines. Further, the number of TAD-like structures is lower in the *syn4* mutants (Fig. 3b; Data S1). Interestingly, the divergence between wild type and *syn4* mutants seems to be more pronounced near chromosome tips (Fig. 3a and 3c). The differences in TAD-like number and position along the chromosome were particularly strong for the third chromosome (Fig. S3). Using TADCompare (Cresswell and Dozmorov, 2020), we found that all known major changes in the TAD structure were observed in the *syn4* mutants at chromosome 3, including new TAD-like structures, changes in their strength, joined or split TAD-like structures (Fig.3d). The TAD-like structures between the mutants are largely comparable (Fig. 3d). From these results we conclude first that SYN4 influences genome structure throughout the *A. thaliana* genome, and that it has a particularly strong influence on inter- and intra-chromosomal interactions in telomere domains.

**Fig. 3.**
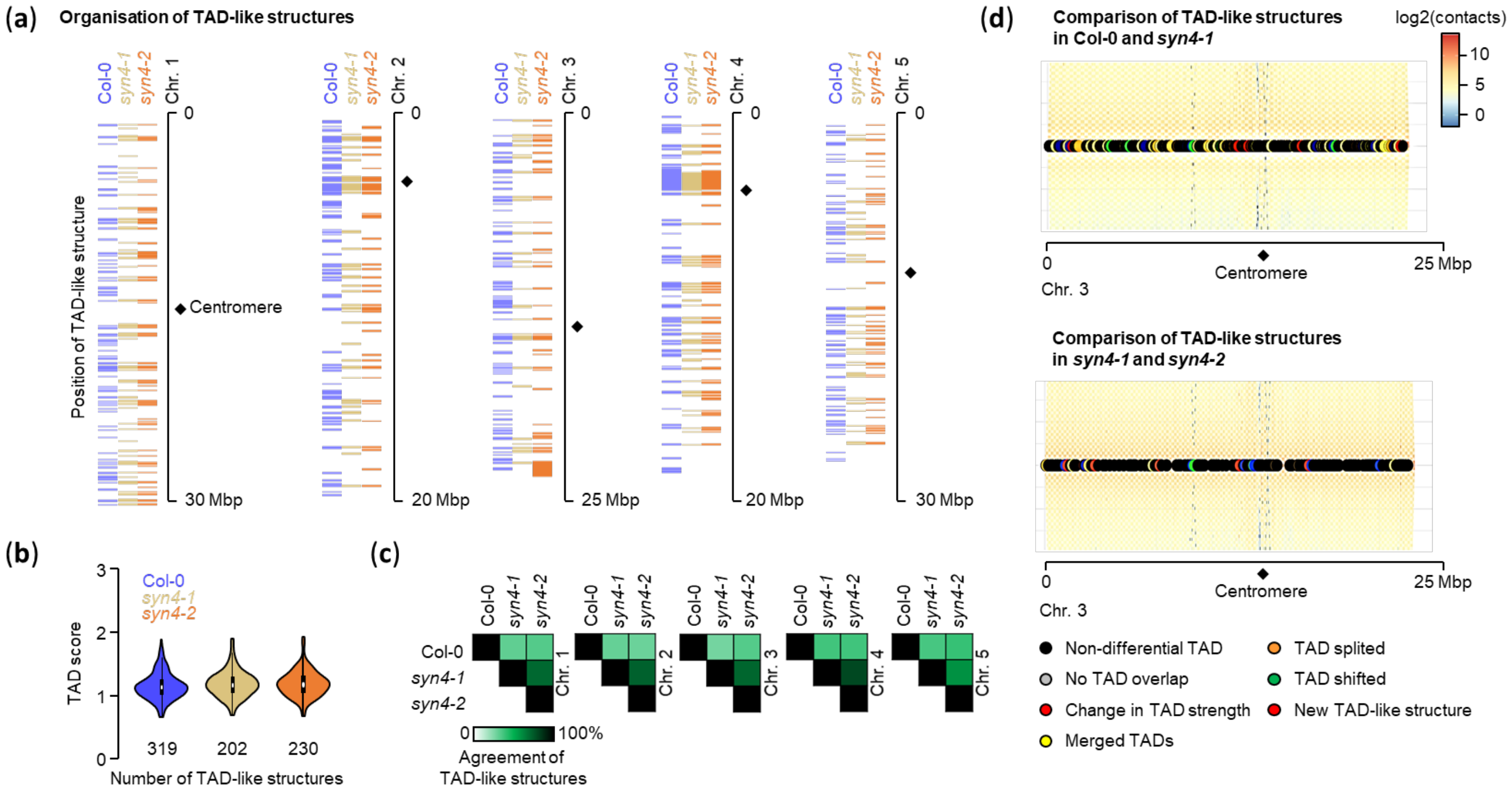
Analyses of TAD-like structures. (a) Altered TAD-like structures, represented by colored bars, are a genome-wide phenomenon in the *syn4*-I and *syn4*-2 α-kleisin mutants compared to Col-0 wild type. (b) No changes in the strength of the TAD-like structures detected, but the number ofT ADs differs between Col-0 wild-type and *syn4* mutants. (d) 61-63% of the wild-type structures cannot be folll.ld in *syn4-l* and *syn4-2*. The TAD agreement in the syn4 mutants is 73% (±5.05%). suggesting that fill ordered fonnation of the genome structure is disrnpted in the absence ofSYN4. (d) TAD-like structures on chromosome 3, which showed the greatest differences in the number of TADs, were compared between Col-0 and *syn4-l* using TADCompare. In *syn4-l* compared to the stmctures in the wild type, strong changes such as shifted, fused and new TADs occur. The mutants among each other show only moderate differences in TAD structure.

### SYN4-dependent genome restructuring affects gene expression

To explore whether the observed altered genome structuring also affects gene expression, we sequenced the transcriptomes of *syn4*-*1* mutant plants and Col-0 wild type. We sampled seedlings grown on culture plates, as well as mature leaves from rosettes. In a partial least squares discriminant analysis (PLS-DA) of transcript profiles from seedlings and leaves from fully grown rosettes (6-week old plants), *syn4-1* gene expression can be clearly distinguished from Col-0 wild type transcripts (Fig. 4a). The number of differentially expressed genes in seedlings is lower than in leaves, 58 and 542 respectively (Fig. 4b; Data S2 and S3). In *syn4-1* seedlings, the expression of *FLOWERING LOCUS T* (*FT*) is reduced 2-fold (Data S2). *FT* expression is shaped by local chromatin loops to which transcription factors such as CONSTANS bind, bringing proximal cis-regulatory elements close to the transcription start site (Cao et al., 2014).

**Fig. 4.**
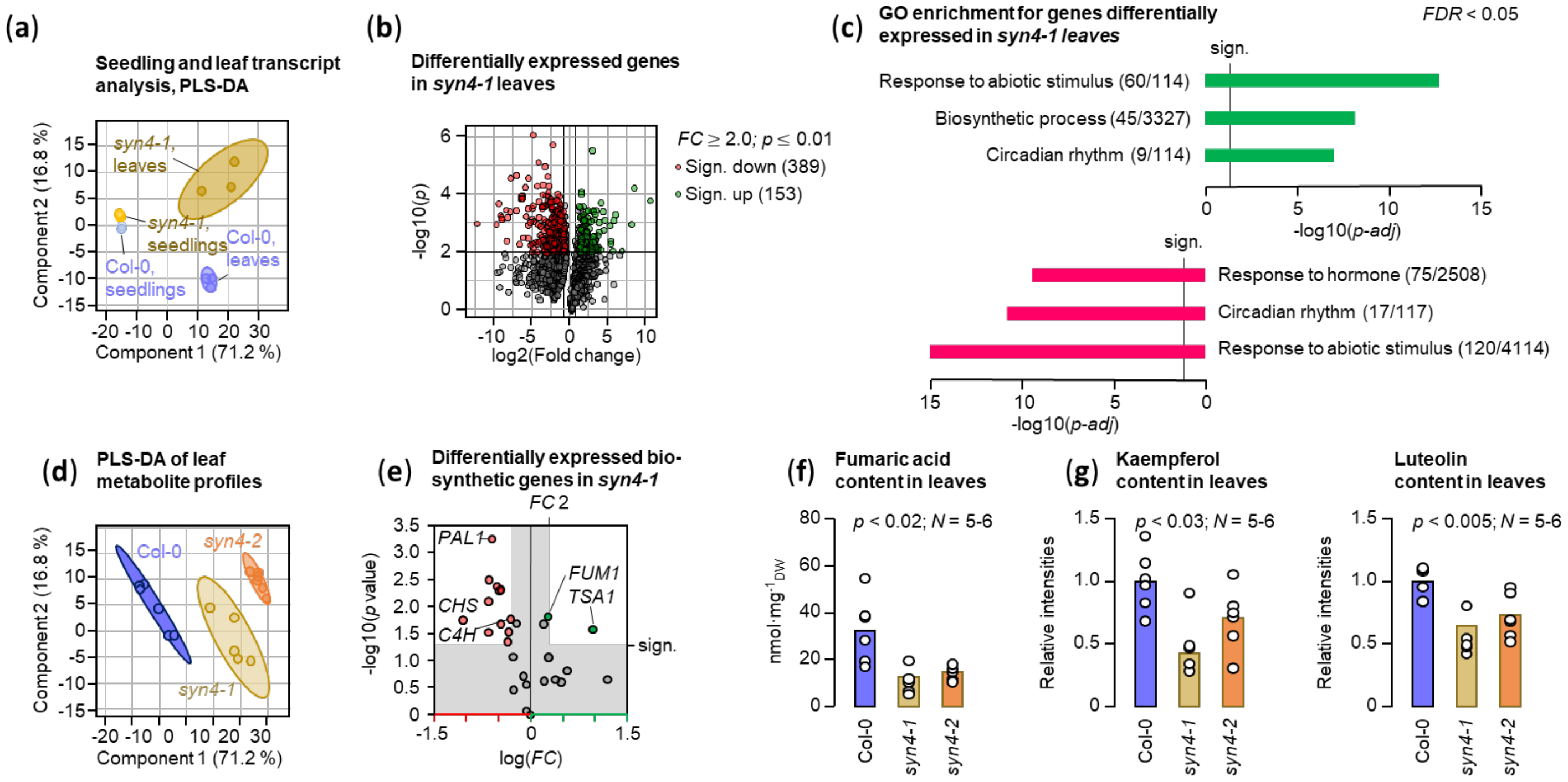
Metabotyping and transcriptional screening of *Arabidopsis* cobesin insertion mutants. (a) PLS-DA of seedling and leaf transcript profiles of the depicted genotypes. The respective 95% confidence interval is shown in the color of the genotype. Tue metabolic profiles of *syn3-I* and *syn4*-I differ most from those of the wild type. (b) The most differentially expressed genes in *syn4*-l relative to Col-0 (*p* ≤ 0.01) from transc1ipt sequencing of mature leaves. (c) GO enrichment analysis (http://geneontology.org/) of up- and down-regulated genes in *syn4*-l leaves showing pathways highly influenced by inse1iion in *SYN4*. For differentially expressed genes in seedlings, GO enrichment analysis did not yield a result. (d) PLS-DA ofleaf metabolite profiles of the Col-0 and *syn4* mutants. The respective 95% confidence interval is shown in the color of the genotype. (e) Biosynthetic genes differentially expressed in Col-0 and *syn4*-l that may have an impact on the metabolome, including *FUMJ* (FUMARASE I), which encodes an enzyme that consumes fumarate in the citrate cycle and produces L-malate. (f) The fumarate content behaves inversely propmiional to the *FUMJ* expression (Student’s T-test between the content of the wild type and the respective *syn4* mutant). (g) Relative kaempferol and luteolin contents in leaves (Student’s T-test between the content of the wild type and the respective *syn4* mutant). The reduced flavonoid content is associated with significantly lower expression in *syn4-1* of early flavonoid biosynthetic genes such as PHENYLALANINE AMMONIA LYASE I (*PALJ*) CINNAMATE-4-HYDROXYLASE (*C4H*) andCHALCONE SYNTHASE (*CHS*).

In leaves, the differentially expressed genes in *syn4-1* cluster more frequently at specific regions of the chromosomes than would occur by chance. This is especially the case for down-regulated genes in *syn4-1* (Data S4; Table S1 and S2; Fig. S4; Cluster locator, http://clusterlocator.bnd.edu.uy/). The observation that seedlings and leaves show very different patterns of differential expression suggests that the role of SYN4 is dynamic and varies among tissues (Fig. 4a). Transcripts of up- and downregulated in *syn4*-*1* leaves are significantly enriched with GO-terms representing specific biological processes (http://geneontology.org/), with transient processes and environmental responses being affected the most (Fig. 4c).

### Metabotyping suggests a physiological consequence of *SYN4* mutation

No distinct visual phenotypes have yet been reported for any of the somatic *Arabidopsis* α-kleisin mutants (Bai et al., 1999; Bhatt et al., 1999; da Costa-Nunes et al., 2014; Dong et al., 2001; Jiang et al., 2007; Yuan et al., 2012). However, because we detected transcriptional alterations, we asked whether these changes might have a less obvious phenotypic output. Therefore, we investigated the “metabotypes” of *syn4* mutants. The “metabotype” or metabolic phenotype provides information about the metabolic state of an organism and is the product of both genetic and environmental factors under specific conditions (Hall et al., 2022; Holmes et al., 2008).

We screened the *syn4* insertion mutants using untargeted LC-MS. Metabolite profiles of *syn4* mutants and Col-0 wild type grown under short-day conditions were obtained from extracts of mature, fully differentiated leaves and compared using PLS-DA to quantify the degree of agreement between the different metabolic fingerprints. We observed substantial differences between metabotypes of wild type Col-0 and *syn4* mutants (Fig. 4d), suggesting altered transcription in these mutants does have a phenotypic consequence for the plant.

We next asked if any of the genes that were transcriptionally altered in the *syn4* mutants could, potentially, directly influence the metabolome (Fig. 4e). The proteins encoded by differentially expressed genes include the enzymes FUMARASE1 (FUM1), and the early flavonoid biosynthetic pathway enzymes PHENYLALANINE AMMONIA LYASE1 (PAL1), CINNAMATE-4-HYDROXLASE (C4H), and CHALCONE SYNTHASE (CHS). When we examined the concentrations of fumaric acid, and the flavonoids kaempferol and luteolin in leaves in more detail, we found that their concentrations were reduced significantly in the mutants (Fig. 4f and 4g). Fumaric acid levels behave inversely to *FUM1* expression, while the flavonoids were positively associated with transcript levels of *PAL1, C4H* and *CHS*, because both transcript levels and the metabolites were reduced in the *syn4* mutants (Fig. 4e-g).

### SYN4 influences the balancing of ABA after induction and drought resilience

The GO-term enrichment analyses of up- and downregulated genes in *syn4-1* suggested that abiotic response processes are influenced by SYN4 function (http://geneontology.org/). We wondered whether water deprivation as an abiotic stress could induce a measurable phenotype. *syn4* mutants reacted less strongly to the lack of water (Fig. 5a). A phytohormone screen revealed that abscisic acid was elevated several fold in the *syn4* mutants after seven days of drought stress (Fig. 5b). We observed a reduction in stomatal conductance with progressive drought in both *syn4* mutant and wild type plants (Fig. S6b). However, the lower stomatal conductance remained stable for several days in the *syn4* mutants (Fig. S6b), whereas stomatal conductance in the wild type decreased further. Furthermore, we noted a more extensive root system in the *syn4* mutants (Fig. S6c-g). Of the wild-type plants, none survived after 14 days of non-watering and re-watering, whereas 14-40% of the mutants recovered (Fig. 5c). To identify the cause of the altered drought phenotype and to reduce the regulatory complexity, the Col-0 wild type and the *syn4-1* mutant were grown on tissue culture plates supplemented with very low ABA concentrations (10 nM). Causal differences were assumed to be present even at low levels of the key phytohormone that regulates the stress response. On control plates no differences in the growth of the seedlings were evident (Fig. 5d). *syn4-1* seedlings were hypersensitive to exogenous treatment with ABA, and showed reduced root growth (Fig. 5d). Numerous ABA hypersensitive mutants are known, but these rarely exhibit increased drought tolerance. Next, the transcriptomes of seedlings grown on 10 nM ABA were obtained. A variety of genes already induced by traces of abscisic acid in the wild type are absent in the *syn4-1* mutant (Data S5). One of the non-induced genes in *syn4-1* is *CYP707A3*, an ABA 8’-hydroxylase that is involved the ABA catabolism (Fig. 5e). The T-DNA insertion mutant *cyp707a3-1* is thought to have higher ABA levels in turgid plants that were hypersensitive to exogenous ABA during early seedling growth (Umezawa et al., 2006). It is further known that the *cyp707a3-1* mutant accumulates a higher amount of stress-induced ABA upon dehydration than the wild type and has a higher drought tolerance (Umezawa et al., 2006). The *cyp707a3-1* drought tolerance phenotype could be reproduced here with both *syn4* mutants in which *CYP707A3* gene expression is deregulated.

**Fig. 5.**
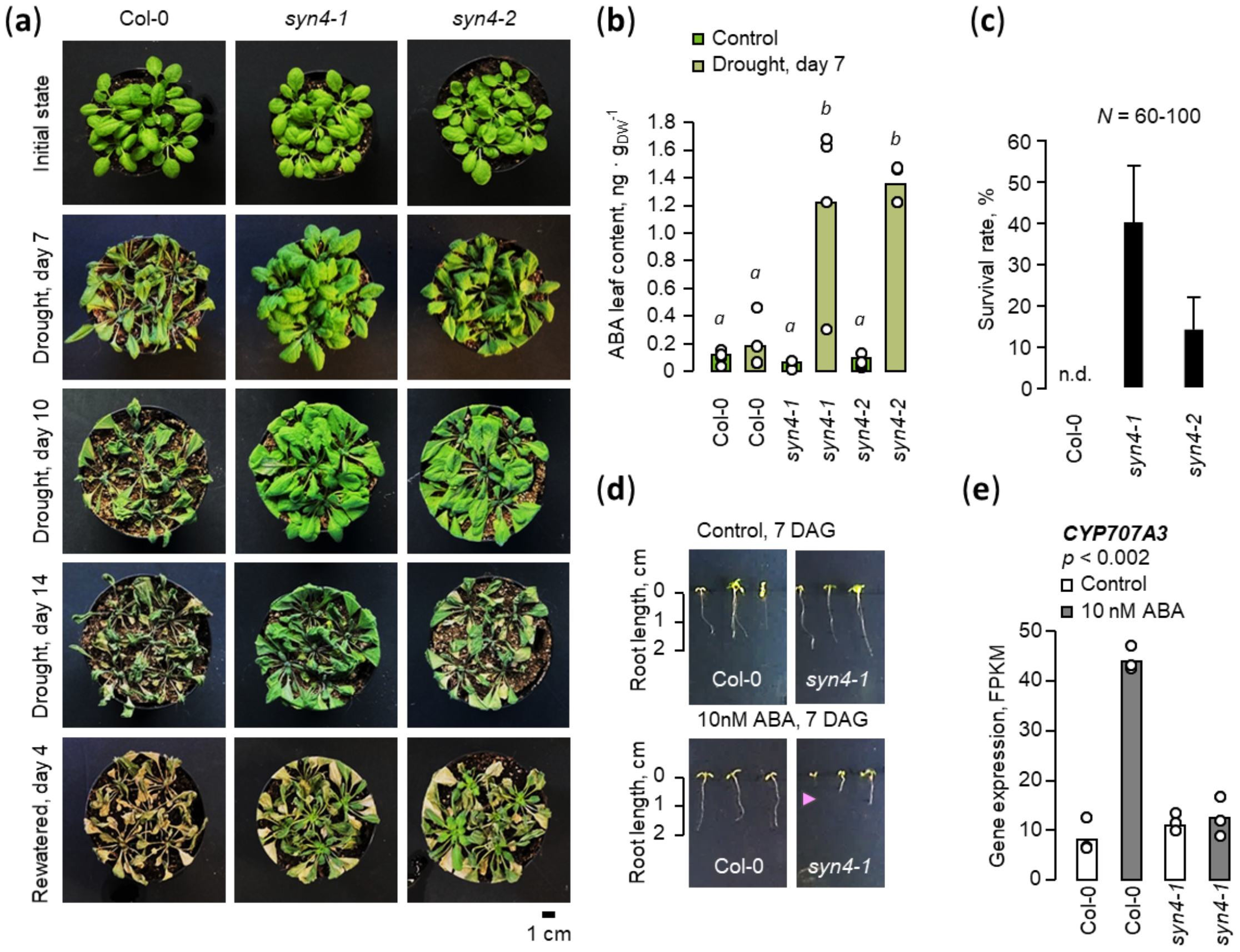
SYN4 influences drought resistance by regulating abscisic acid contents. (a) Plants were not watered for 14 days and photographed on the days indicated after the stmt of the water withdrawal. The photos are representative for entire batches. The expennint comprised 60-100 plants for each genotype. (b) Abscisic acid concentrations me higher in the *syn4* mutants than in the wild type after seven days of drought stress. (One-way ANOVA; *N* = 4). (c) The SU1vival rate after re-watering was detennined. No plants were found to survive the ch-ought stress for the Col-0 wild type. *syn4* mutants, on the contraiy, show a survival rate of 40% and 14% for *syn4*-l and *syn4*-2, respectively. (d) *syn4* a-kleisin mutants on culture plates are hypersensitive to low concentrations of abscisic acid (10 nM ABA) under long-day conditions. (e) CYP707A3, which encodes an ABA 8’-hydroxylase involved in ABA catabolism, is not induced upon treatment with 10 nM ABA in *syn4*-l mutants (Student’s t-test, *N* = 3).

We examined the extent to which the chromatin structure at the *CYP707A3* locus differs in the wild type and *syn4* mutants. In untreated leaves, TAD-like structures around the *CYP707A3* locus differ between the wild type and *syn4* mutants (Fig. 6a and 6b). In the wild type, the locus is located at a TAD boundary, whereas in the mutants the locus is located within the shifted TAD-like structure (Fig. 6a and 6b). Directly at the locus, enhanced DNA interactions in the *CYP707A3* promoter region and within the gene body were detected in the mutants (Fig. 6c) which might affect functional enhancer-promoter interaction (EPIs; Fig. 6c, wild type loop no. 1). Enhancer interactions are still detectable after 7 days of drought stress, determined using FRET probes (Fig. 6d). In general, dis-functional DNA interactions at the promoter site can disrupt promoter activity and thus affect induced gene expression, leading to an overall altered stress response (Fig. S11a; Furlong and Levine, 2018).

**Fig. 6.**
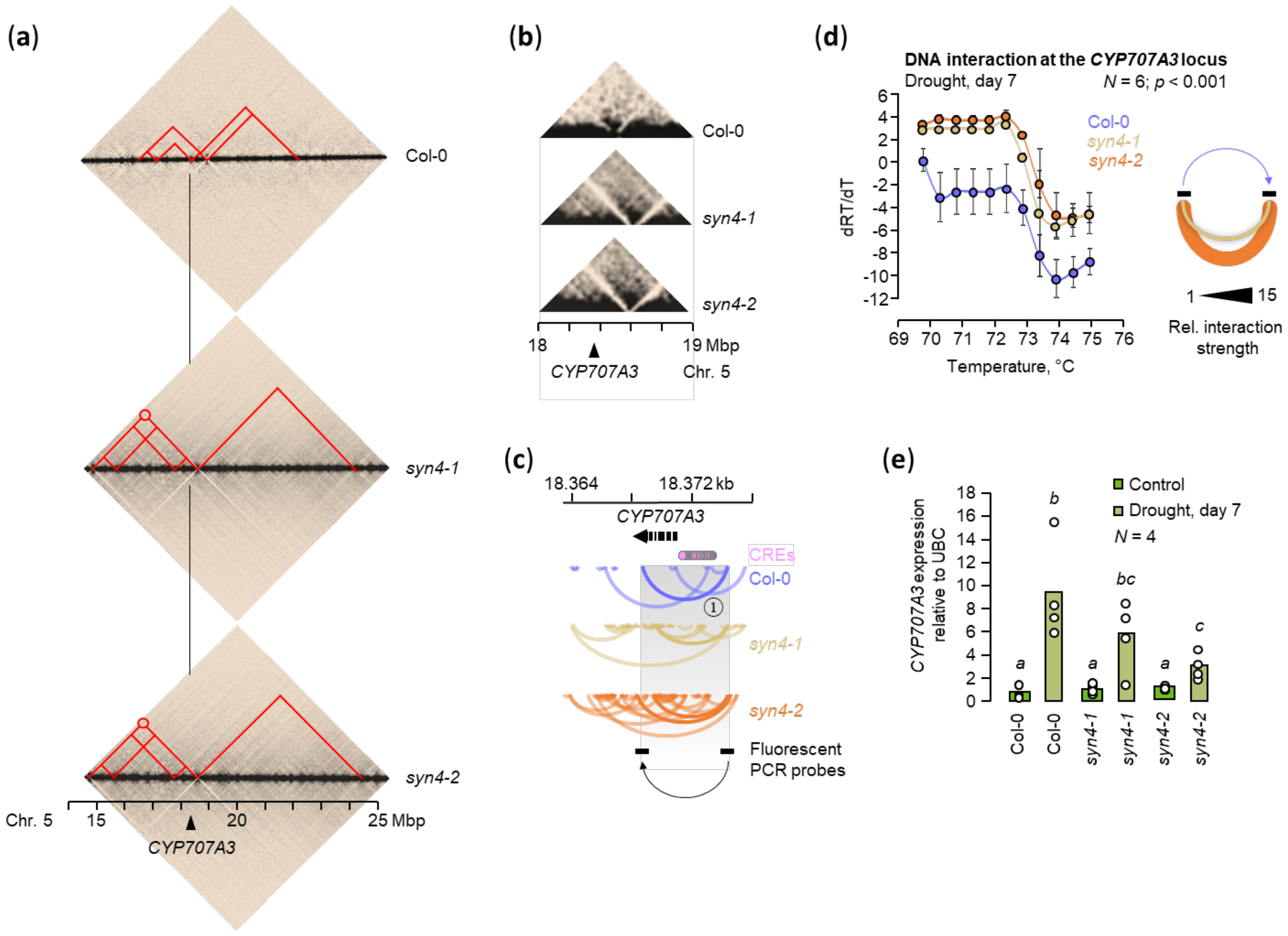
Influence of SYN4 on chromatin architecture at the *CYP707A3* locus and its consequences on gene expression under water deprivation. (a and b) Hi-C interactions near the *CYP707A3* locus generated :from Col-0, *syn4*-l and *syn4*-2 merged leaf samples. The heat maps were displayed with Juicebox I. I I.OS and the maps were normalized based on the coverage (Sqrt). The resolution was 25 kbp and the respective interactions were displayed in grey values. (c) DNA interactions at fue *CYP707A3* locus :from un1reated Col-0, *syn4*-I and *syn4*-2 leaves. Cis-regulatory element (CRE) binding sites were predicted using AGRIS (https://agris-knowledgebase.org/). Fluorescent probes were developed for the *CYP707A3* promoter region and the gene body spanned by loop no. I to detennine whether and to what extent these chromatin regions interact with each other. The binding sites are labeled. (d) Change in fluorescence over time signalling chromatin interactions at fue *CYP707A3* locus for plants after 7 days of water deprivation. As under control conditions, chromatin interactions are strongly increased in the *syn4* mutants under drought stress. (e) *CYP707.4.3* gene expression in drought-stressed plants (One-way ANOVA; *N* = 4).

### SYN4 is conserved throughout land plants

The data presented suggest that SYN4, like the other plant α-kleisins proteins, has a specific cellular function, which may be due to its distinct protein structure. The modelled secondary structures (http://bioinf.cs.ucl.ac.uk/psipred) of all SYN proteins indicate that they have in common globular helical domains at the N- and C-termini, which serve the interaction with the SMC arms (Fig. S 5a). This is also consistent with the characterized human α-kleisins REC8 and SCC1 / RAD21. The main difference, however, is the length of the unstructured areas in SYN4, which exceeds that of the characterized eukaryotic α-kleisins. This unique structural feature of SYN4 may indicate a function not found in other eukaryotes.

Since no clear homolog of SYN4 is found in fungi or animals (Fig. S 5b), we asked if it is found in other plants. Therefore, we performed a phylogenetic analysis of eukaryotic α-kleisins together with the plant sequences (Data S6; https://www.ncbi.nlm.nih.gov/gene). All four RAD21-like α-kleisin genes occur across land plants, suggesting their duplication is ancient, and not unique to *A. thaliana* (Fig. S5b).

## Discussion

Plants have at least four unique cohesin complexes due to the presence of four distinct α-kleisins. Their known roles include conserved eukaryotic functions in sister chromatid cohesion, meiotic axis formation and recombination, and double strand break repair (Bai et al., 1999; Bhatt et al., 1999; Bolaños-Villegas et al., 2017; da Costa-Nunes et al., 2006; Jiang et al., 2007). However, whether the three plant cohesin complexes expressed in mitotic cells in *A. thaliana* (complexes containing the α-kleisins SYN2 - 4) also affect chromatin structure, as is the case in other eukaryotes, has not yet been investigated. Here we show in *A. thaliana* that SYN4, a somatic α-kleisin conserved in all plants, indeed affects genome structure, most notably telemore interactions, gene expression and metabolite biosynthesis in somatic cells.

It is established that eukaryotic genomes are organized hierarchically (Kim et al., 2019; Davidson et al., 2019). Chromatin loops form the basis for larger-scale topologically-associated-domains (TADs) that are associated with particular epigenetic marks and play an important role in gene regulation (Szabo et al., 2019). Our data show that SYN4-containing cohesins contribute to TAD-like domain formation in *A. thaliana*. SYN4 is, however, only one of three somatic α-kleisins in *A. thaliana* (the other two being SYN2 and SYN3) and it is possible they all play complementary and perhaps partially overlapping roles in genome organization.

The loss of SYN4 activity has a pronounced effect on telomere interactions among chromosomes. Telomere interactions and associated effects on gene expression in somatic cells are observed across eukaryotes (Robin et al., 2014; Taddei et al., 2009), and telomere clustering has been reported from Hi-C as well as cytological studies in a wide range of plants including *A. thaliana* (Armstrong et al., 2001; Feng et al., 2014; Fransz et al., 2002; Moissiard et al., 2012; Schubert et al., 2012;). The role of somatic telomere clustering is not well understood, although it might relate to the assembly of so-called “transcription factories” (Xu and Cook, 2008). It has been suggested that DNA-DNA interaction of repetitive elements such as telomere sequence motifs might be needed for the formation these transcription factories (Xu and Cook, 2008). Therefore, it would be interesting in future to investigate whether the effect of SYN4 (and/or the other “mitotic” α-kleisins in plants) on telomere interactions influences the assembly of somatic transcription factories and/or the formation of meiotic telomere bouquets, as SYN1/REC8 does during meiosis (Trelles-Sticken et al., 2005).

How somatic telomere clustering in plants comes about at the molecular level is even less well understood than its role. The fact that the loss of SYN4 almost completely disrupts interchromosomal telomere interactions is remarkable because we know of no other mutant in plants that has this effect in somatic cells. The nature of the connection between TAD-like domains and histone modifications, and whether the effect of SYN4 is direct or indirect, is still unclear. However, the fact that the *syn4* mutant so strongly affects these interactions is a valuable starting point to better understand the molecular coordination of DNA architecture in eukaryotic cells.

We observed that the *syn4* mutants show altered transcript levels (Fig. 4a and 4b), suggesting the alterations to chromatin organization have detectable consequences for the plant. This is further supported by the observation that the metabolome is substantially affected (Fig. 4d). Transcriptional changes in *syn4* mutants suggest that the effect of SYN4 might largely be on genes involved in environmental responses (Fig. 4c). Using the *CYP707A3* locus as an example, we found that the existing genome architecture has an influence on the induction of gene expression. It is well established that the relative positioning of loops to *cis* regulatory elements (CREs) and the promoter have a decisive influence on functional enhancer-promoter interactions und thus gene expression (Fig. S10a; Furlong and Levine, 2018). If this is changed, as here for the *CYP707A3* locus, it can have an influence on the induction of gene expression. This could explain why genome structures differ greatly between plants, unlike e.g. in vertebrates (Szabo et al., 2019), because each plant species may have adapted its genome structure and EPIs (positioning of CREs) to the respective location with the specific given habitats. We propose that SYN4 is involved in this adaptation by controlling the distribution of loops and higher chromatin architecture (Fig. S10b). We hypothesize that SYN4 is involved in this adaptation by controlling the distribution of loops and higher chromatin architecture (Fig. S11b), which presumably depends on the plant species and its ecosystem and leads to a specific and unique plant genome structure.

## Acknowledgments

This research was funded by UK BBSRC (Institute Strategic Programme GEN BB/P013511/1, AND), The Leverhulme Trust (RPG-2018-216, AND), DFG (German Research Foundation, Walter Benjamin Program HA8855/1-1, VH) and by core funding to the Leibniz Institute of Plant Biochemistry (TR).

## Author Contributions

Conceptualization: PP, TR, KB, AND, VH

Methodology: PP, VH

Investigation: PP, CP, SK, KG, JZ, VH

Visualization: PP and VH

Supervision: TR, VH

Writing—original draft: PP, VH

Writing—review & editing: TR, AND, KB, VH

All authors read and approved the final manuscript.

## Data availability

The RNASeq and Hi-C data for this study are freely available in the European Nucleotide Archive (ENA; https://www.ebi.ac.uk/ena) and will be available upon publication with the project ID PRJEB53624.

