## Supplemental File 1 for "The plant specific cohesin subunit SYN4 contributes to 3D genome organization"

Pirita Paajanen *et al.*

**This PDF file includes:**

Figs. S1 to S10
Tables S1 to S2

**Other Supplementary Materials for this manuscript include the following:**

Data S1 to S6

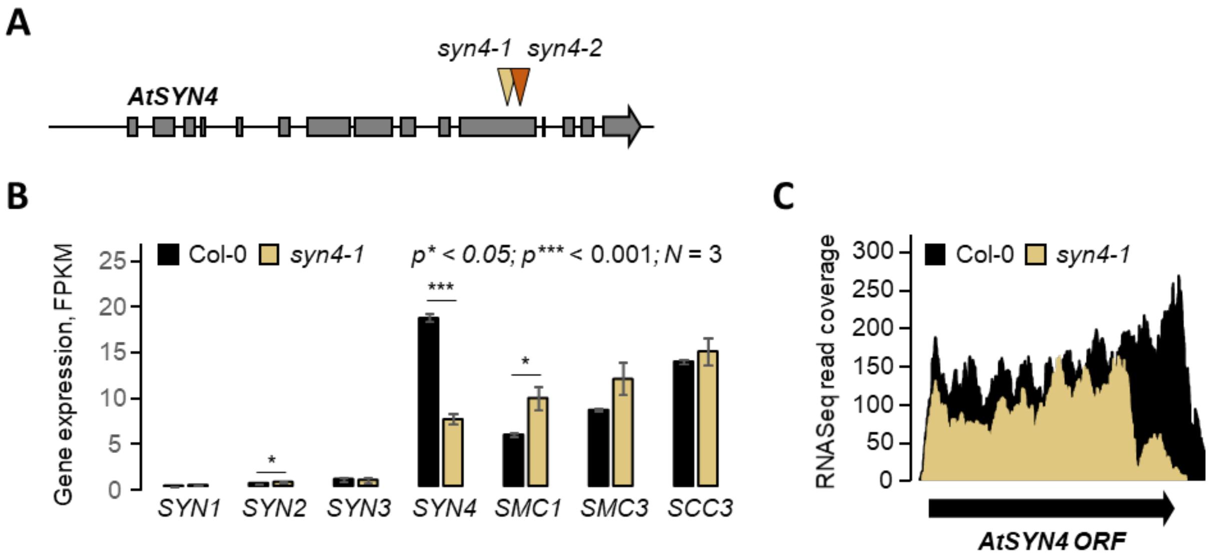


**Fig. S1. Expression of cohesin genes in Col-0 and *syn4-1* mature leaf tissues.**

(A) Intron-exon map of the *AtSYN4* locus in Arabidopsis thaliana. The T-DNA insertion sites for the stable homozygous lines *syn4-1* (SALK_076116) and *syn4-2* (SALK_020171) are marked in ochre and orange, respectively.

(B) Expression of cohesin genes in mature leaves of the Col-0 wild type and the *syn4-1* mutant. *SYN4* expression is strongly decreased in the mutant (paire-wise Student's T-test between Col-0 and *syn4-1* measurements, bars represent standard errors, s.e.).

(C) RNA sequencing reads for the *AtSYN4* locus from the wild-type, Col-0, and the *syn4-1* mutant show that only a truncated non-functional *SYN4* coding sequence is expressed in *syn4-1*.

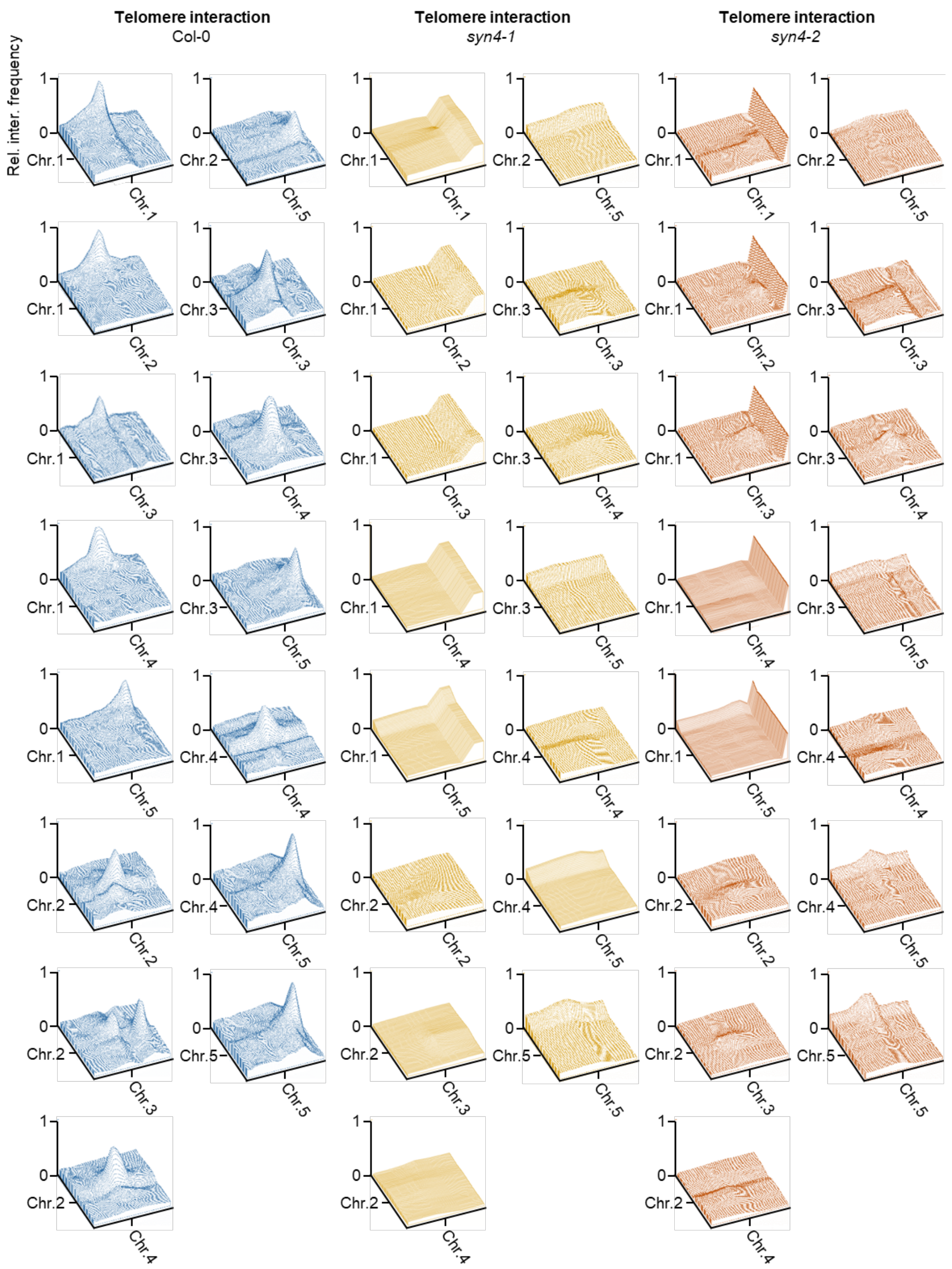


**Fig. S2. Chromatin interaction analysis.**

Surface plots show interactions of telomeres generated from Hi-C heat maps using imageJ. The heat maps were displayed with Juicebox 1.11.08 and the maps were normalized based on the coverage (Sqrt). The relative interaction frequency always refers to the strongest blackening in the respective map.

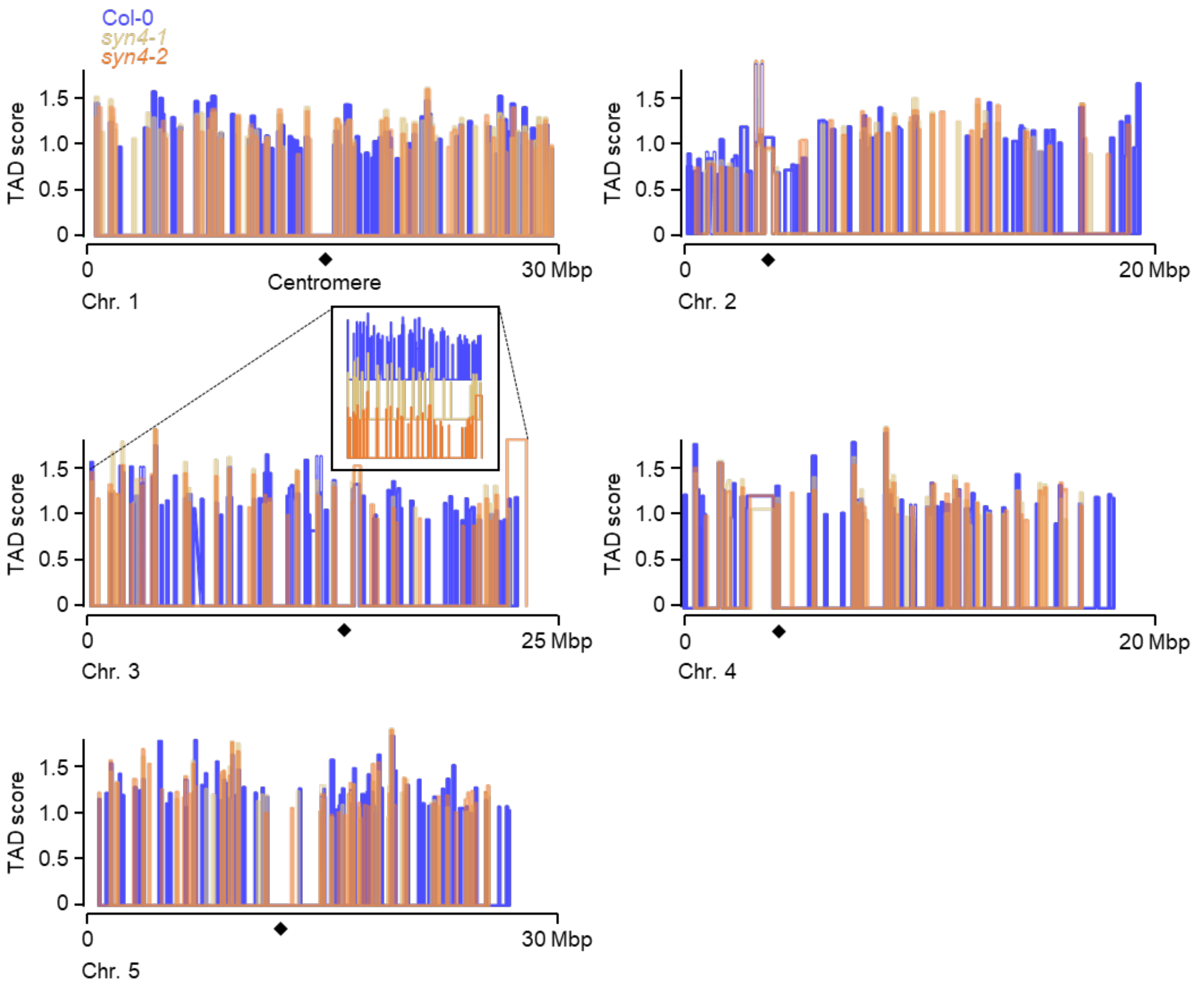


**Fig. S3. TAD-like structure and distribution analysis of the individual *Arabidopsis* chromosomes.**

The position and strength of the TAD-like structures were plotted along the chromosomes. For chromosome 3, only 42-59% of the number of wild-type TADs occur in the mutants.

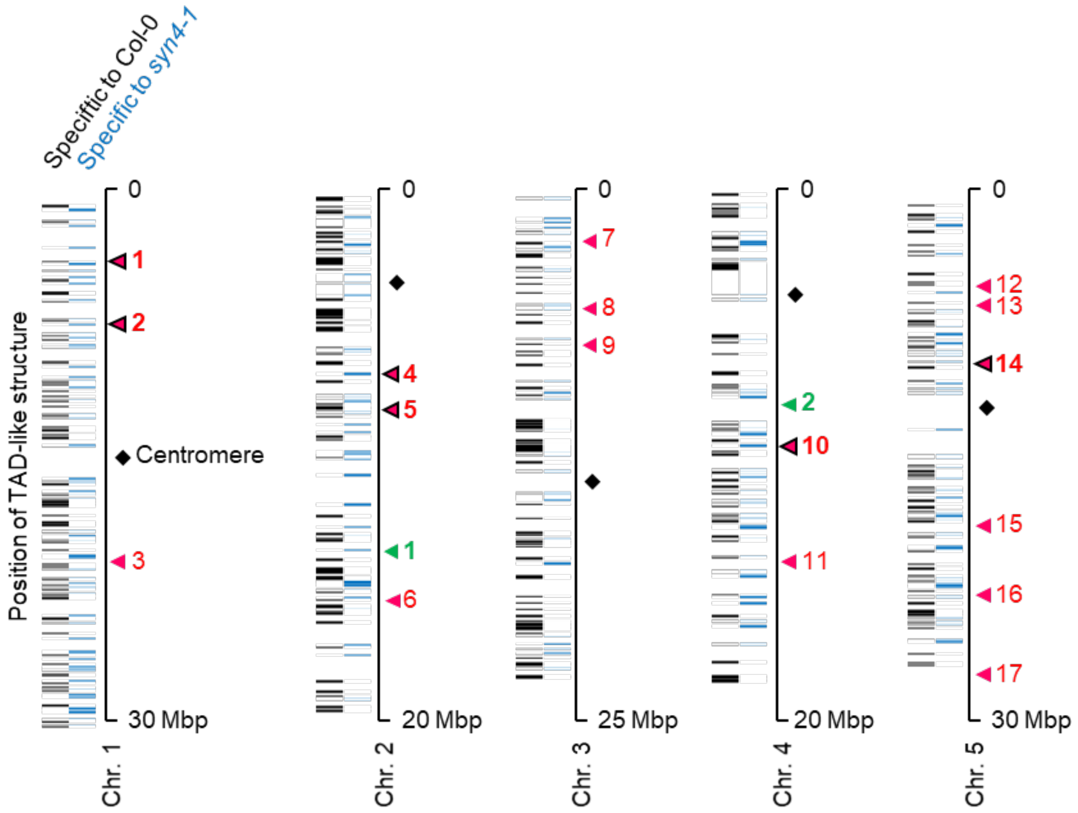


**Fig. S4. Specificity of the TAD regions for Col-0 and *syn4-1*.**

TAD regions that only occur in the wild type are shown in black and those that are specific to *syn4-1* are depicted in blue. The arrows indicate co-regulated gene clusters. Clusters whose genes in *syn4-1* show low expression are marked in red. Clusters with genes that are upregulated are marked in green. Some of the gene clusters are located in regions with private TADs (arrows outlined in black).

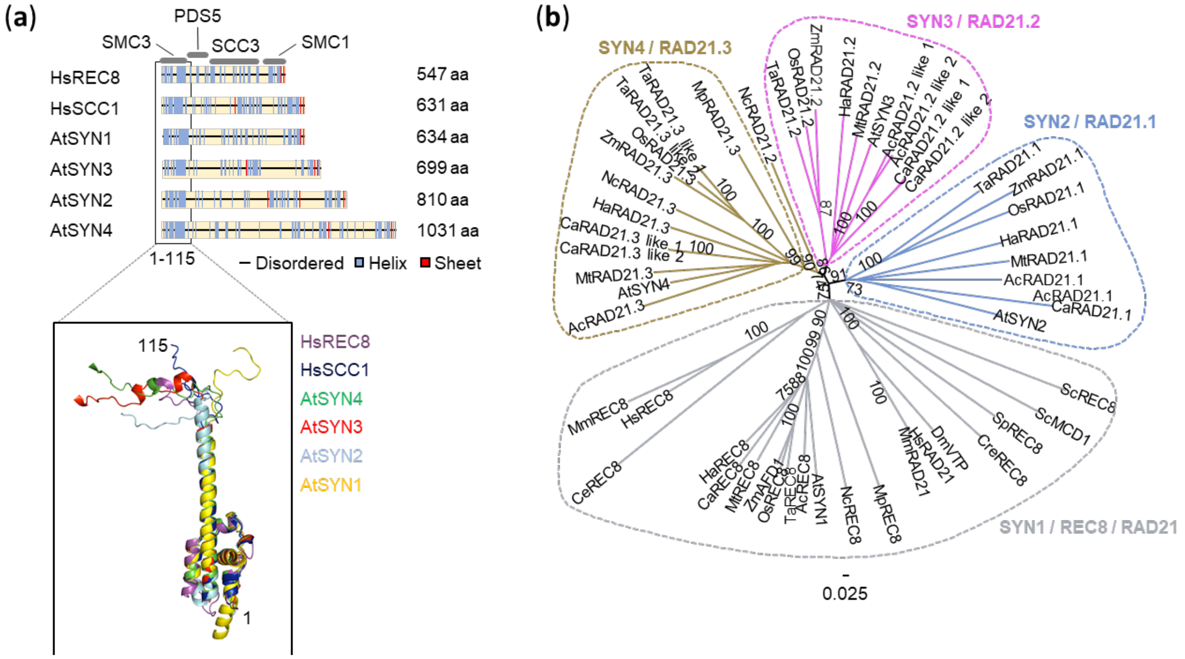


**Fig. S5. α-kleisin structure conservation.**

(a) Secondary protein structures of individual α-kleisins from humans (Hs) and *Arabidopsis thaliana* (At). The amino acid sequence of the individual proteins are highly variable (Identity < 17%). The secondary structure shows some common motifs, such as the helical SMC binding regions. The secondary protein structures of HsREC8 and HsSCC1, α-kleisin proteins known to function during meiosis and mitosis in humans, respectively, and the interaction sites with associated proteins are shown as references. The window shows the modelled structures (https://alphafold.ebi.ac.uk/) of the N-terminus (1-115aa) of the investigated α-kleisins. The structures were aligned with the PyMOL super-alignment function.

(b) Unrooted phylogeny of eukaryotic α-kleisin protein sequences. Annotated sequences for *Arabidopsis thaliana* (At), *Caenorhabditis elegans* (Ce), *Drosophila melanogaster* (Dm), *Helianthus annuus* (Ha), *Homo sapiens* (Hs), *Medicago truncatula* (Mt), *Mus musculus* (Mm), *Oryza sativa* (Os), *Saccharomyces cerevisiae* (Sc), *Schizosaccharomyces pombe* (Sp), *Triticum aestivum* (Ta) and *Zea mays* (Zm) were downloaded from the ncbi website (https://www.ncbi.nlm.nih.gov/gene). For a BLAST search, the conserved N-terminus of a species of the next higher plant class was used to identify α-kleisin sequences in these species: *Aquilegia coerulea* (Ac), *Chlamydomonas reinhardtii* (Cre), *Coffea arabica* (Ca), *Marchantia polymorpha* (Mp), *Nymphaea colorata* (Nc). The protein names are preceded by the species as abbreviation. The bootstrap (>75) are given at the branches.

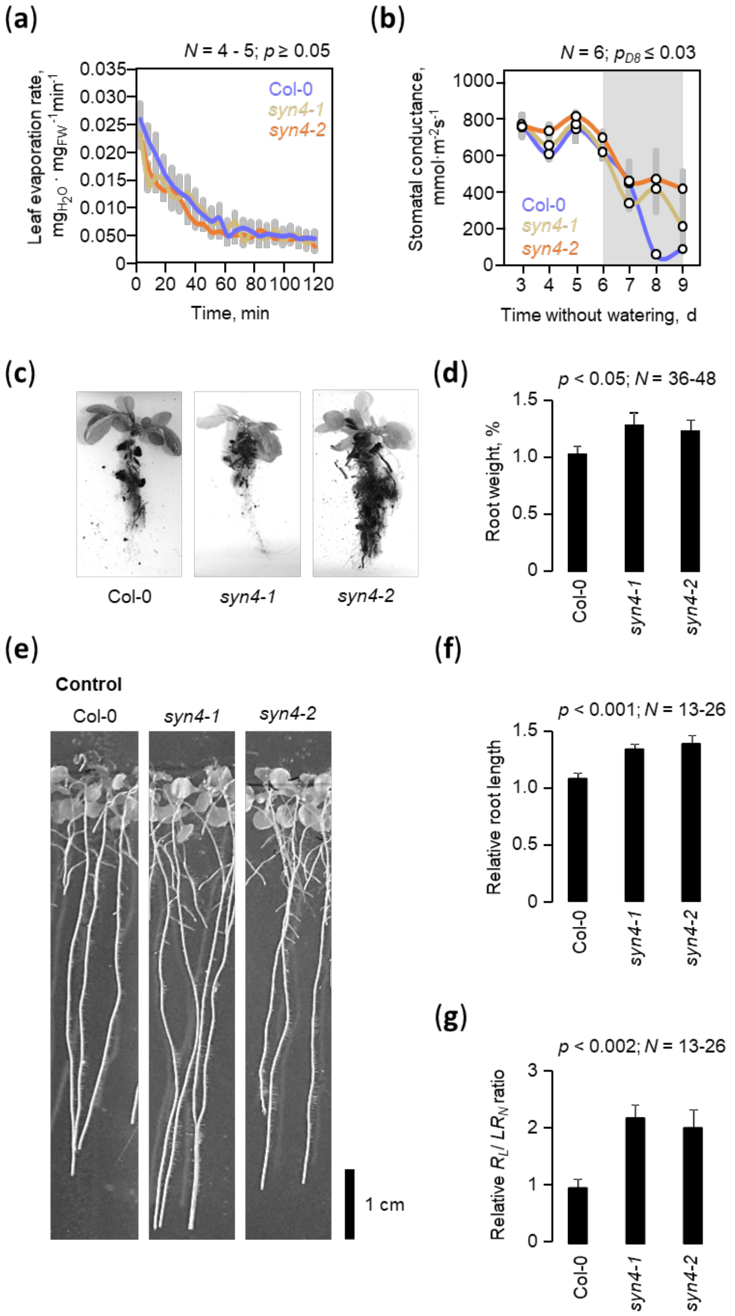


**Fig. S6. SYN4 influences drought resistance by regulating abscisic acid contents.**
(a) Leaf evaporation rate of leaves of untreated plants over two hours. No significant differences were found.
(b) Stomatal conductance between day 3 and day 9 after omitted watering ($N = 6$). Leaf conductance decreases later in the *syn4* mutants than in the Col-0 wild type (Student's t-test, s.e.).
(c) The roots of 6-week-old *Arabidopsis* plants suggested a morpholigical difference in *syn4* mutants.
(d) *syn4* plants in hydroponic cultures develop more roots (Student's t-test; s.e.)
(e-g) 14d old *syn4* seedling on culture plates produce longer roots ($R_L$) and more side roots ($R_N$) relative to Col-0 wild type (Student's t-test; s.e.).

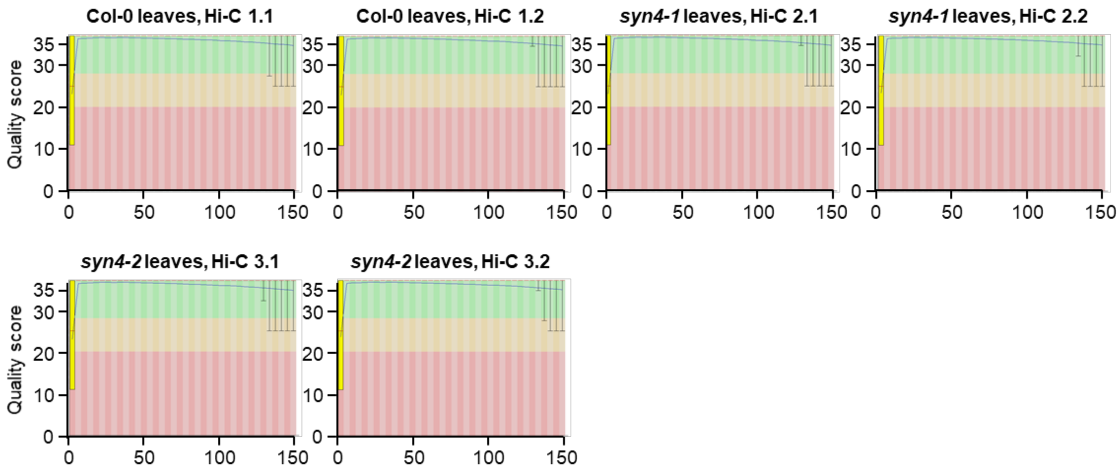


**Fig. S7. Per Base Sequence Quality analysis of Illumina sequences from *Arabidopsis* leaf Hi-C libraries .** Overview of the range of quality values across all bases at each position in the FastQ file. The blue line represents the mean quality and central red line is the median value. Paired-end 150 bp sequences are from Col-0 and *syn4-1, syn4-2* Hi-C samples (*N* = 1).

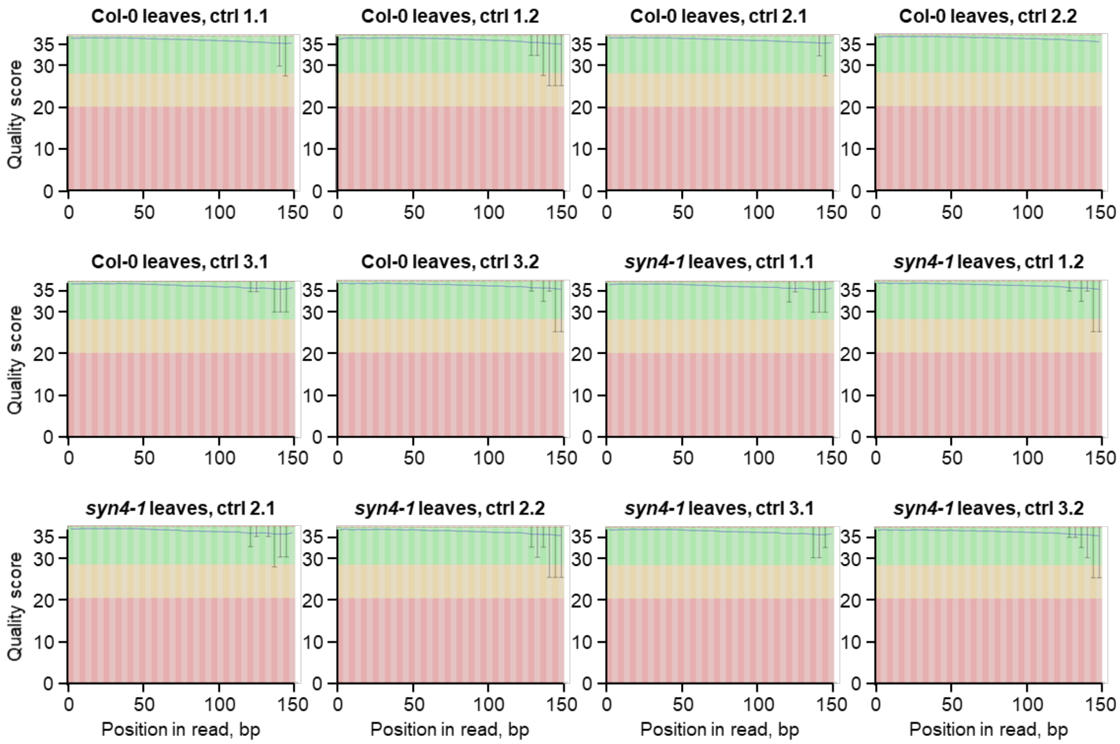


**Fig. S8. Per Base Sequence Quality analysis of Illumina sequences from RNA leaf samples.** Overview of the range of quality values across all bases at each position in the FastQ file. The blue line represents the mean quality and central red line is the median value. Paired-end 150 bp sequences are from Col-0 and *syn4-1* RNA leaf samples ($N = 3$).

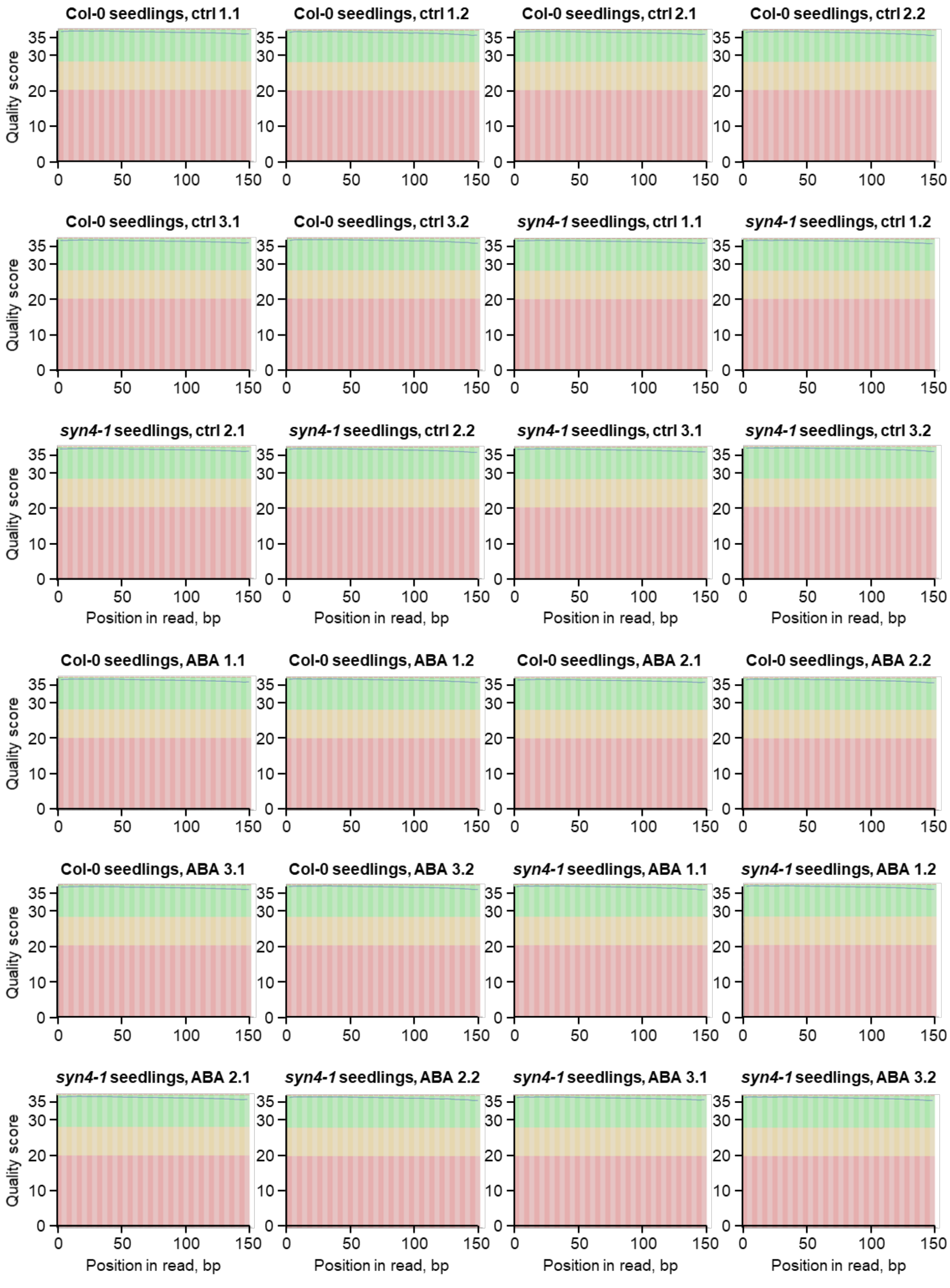


**Fig. S9. Per Base Sequence Quality analysis of Illumina sequences from RNA seedling samples.** Overview of the range of quality values across all bases at each position in the FastQ file. The blue line represents the mean quality and central red line is the median value. Paired-end 150 bp sequences are from Col-0 and *syn4-1* RNA samples with and without the addition of 10nM abscisic acid (ABA; N = 3).

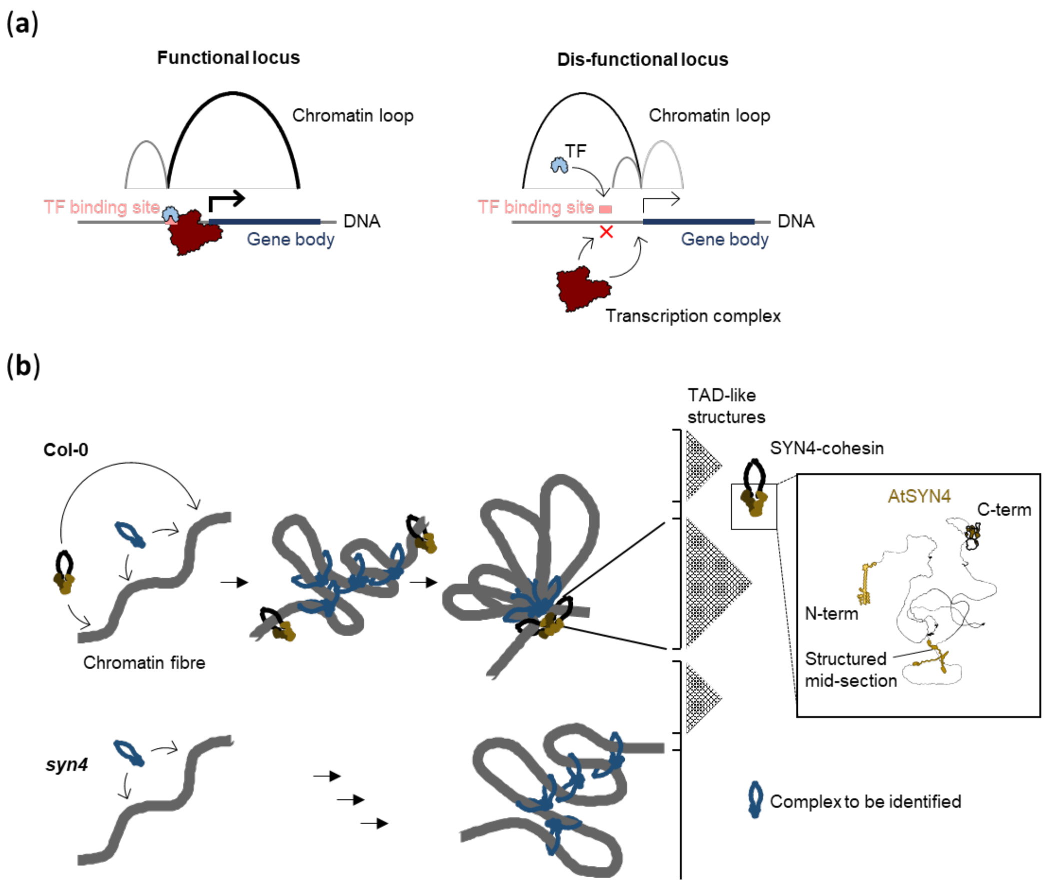


**Fig. S10. Models for SYN4 functioning.** (**a**) The relative positioning of chromatin loops is crucial for the interaction of enhancer and promoter. When physically separated by loops, contact between molecular factors in the enhancer and promoter regions is reduced, resulting in decreased gene expression (dis-functional locus). Expression of genes can occur if these isolating chromatin structures are not present (functional locus). (**b**) SYN4-cohesins influence higher genome structures and the positioning of loops. We postulate that SYN4-cohesins stabilize TAD-like structures by binding two distant chromatin regions with the SMC arms and through SYN4. Within these by SYN4 restricted TAD-like structures, loops are formed by as yet undetermined factors, possibly other cohesin complexes. In the *syn4* insertion mutants, these TAD-like structures would be absent and the loops could be formed beyond the TAD boundaries. The window shows a AtSYN4 protein model generated by AlphaFold (https://alphafold.ebi.ac.uk/) and graphically processed using Inkscape. The structured areas are shown in color.

**Table S1. Clustering in differentially expressed genes from *syn4-1* seedlings[1].**

| | $C_{Random}$ | s.d. | $C_{Leave\ tissues}$ | Number | *p*-value |
|---|---|---|---|---|---|
| Up-reg. | 0.09% | ±1.19% | 0% | 0 | 4.69e-01 |
| Down-reg. | 0.34% | ±1.23% | 0% | 0 | 3.90e-01 |

[1]http://clusterlocator.bnd.edu.uy/; Max-gap = 0

**Table S2. Clustering in differentially expressed genes from *syn4-1* leaf tissues[1].**

| | $C_{Random}$ | s.d. | $C_{Leave\ tissues}$ | Number | *p*-value |
|---|---|---|---|---|---|
| Up-reg. | 1.1% | ±1.18% | 3.33% | 2 | 2.88e-02 |
| Down-reg. | 2.78% | ±1.19% | 9.07% | 17 | 6.02e-08 |

[1]http://clusterlocator.bnd.edu.uy/; Max-gap = 0
